# Axonal growth of cortical neurons does not require paxillin

**DOI:** 10.64898/2025.12.11.693761

**Authors:** Katelyn Rygel, Kasey Gillespie, Kristy Welshhans

## Abstract

Axon growth is an essential cellular process during neural development, and its dysregulation contributes to numerous neurodevelopmental disorders. During axon growth, extracellular signals direct neurons to extend projections that connect with their synaptic targets. Paxillin is a key member of adhesion sites that control motility by linking the intracellular actin cytoskeleton to the extracellular matrix. Paxillin also binds to the cytoskeletal protein, tubulin. However, little is known about the role of adhesion proteins in neurons. Here, we use conditional paxillin knockout mice to investigate how loss of paxillin in pyramidal cortical neurons affects developing neuron morphology. Surprisingly, loss of paxillin in pyramidal cortical neurons caused no change in axon length or soma area between control (*Pxn*^*F/F*^) and conditional paxillin knockout (*Pxn*^*F/F; Emx1-Cre*^) mice at basal conditions. Following brain-derived neurotrophic factor stimulation, the loss of paxillin resulted in no change in soma area or axonal β-tubulin levels, but did result in a significant increase in axon length, as compared to control. Finally, the corpus callosum size was not significantly different between *Pxn*^*F/F*^ and *Pxn*^*F/F; Emx1-Cre*^ animals. In summary, these data suggest that paxillin is not required for axonal growth during neural development.

## 1. INTRODUCTION

During neural development, neurons extend axons that connect with their synaptic targets. One cellular mechanism regulating axon extension involves adhesion complexes, which link the extracellular matrix to the intracellular actin cytoskeleton, restricting the retrograde flow of actin and resulting in membrane protrusion^1^. Adhesion sites contain many proteins, including paxillin, which is a scaffolding protein. Previous work investigating paxillin loss in neural progenitor cells (NPCs) showed that these cells migrate more slowly than controls, resulting in a delay in cortical layer formation^2^.

Tubulin is a heterodimer protein that assembles into microtubules, a primary cytoskeletal component of the axon shaft ^3^. Interestingly, tubulin binds to the LIM2-LIM3 domains of paxillin. Furthermore, paxillin has been shown to regulate microtubule dynamics in non-neuronal cells by controlling catastrophe, the rapid disassembly of microtubules^4^. Here, we use conditional paxillin knockout mice to examine neuron morphology and expression of β-tubulin in developing pyramidal cortical neurons that do not express paxillin.

## 2. RESULTS AND DISCUSSION

Global knockout of paxillin is embryonic lethal by embryonic day 9.5 (E9.5)^5^; therefore, we used paxillin floxed animals (*Pxn*^*F/F*^) crossed with Emx-1 Cre mice to generate conditional paxillin knockout mice (*Pxn*^*F/F; Emx1-Cre*^). These mice have paxillin knocked out in ∼88% of pyramidal neurons in the developing cortex and hippocampus^2, 6^. We extracted cortex from control *(Pxn*^*F/F*^) and conditional paxillin knockout (*Pxn*^*F/F; Emx1-Cre*^) animals at E17, and validated paxillin knockout by digital droplet PCR (RT-ddPCR) and immunocytochemistry (**Figure 1A-B**). RT-ddPCR showed a reduction in paxillin in *Pxn*^*F/F; Emx1-Cre*^ mice that approached significance (p = 0.0571), but this experiment used whole cortex, which has a mixed population of cell types, some of which are not pyramidal neurons and thus still contained paxillin (**Figure 1A**). Using quantitative immunocytochemistry to quantify paxillin only in cortical pyramidal neurons, there was a significant reduction in paxillin in neurons from *Pxn*^*F/F; Emx1-Cre*^ embryos (**Figure 1B**).

**Figure 1.**
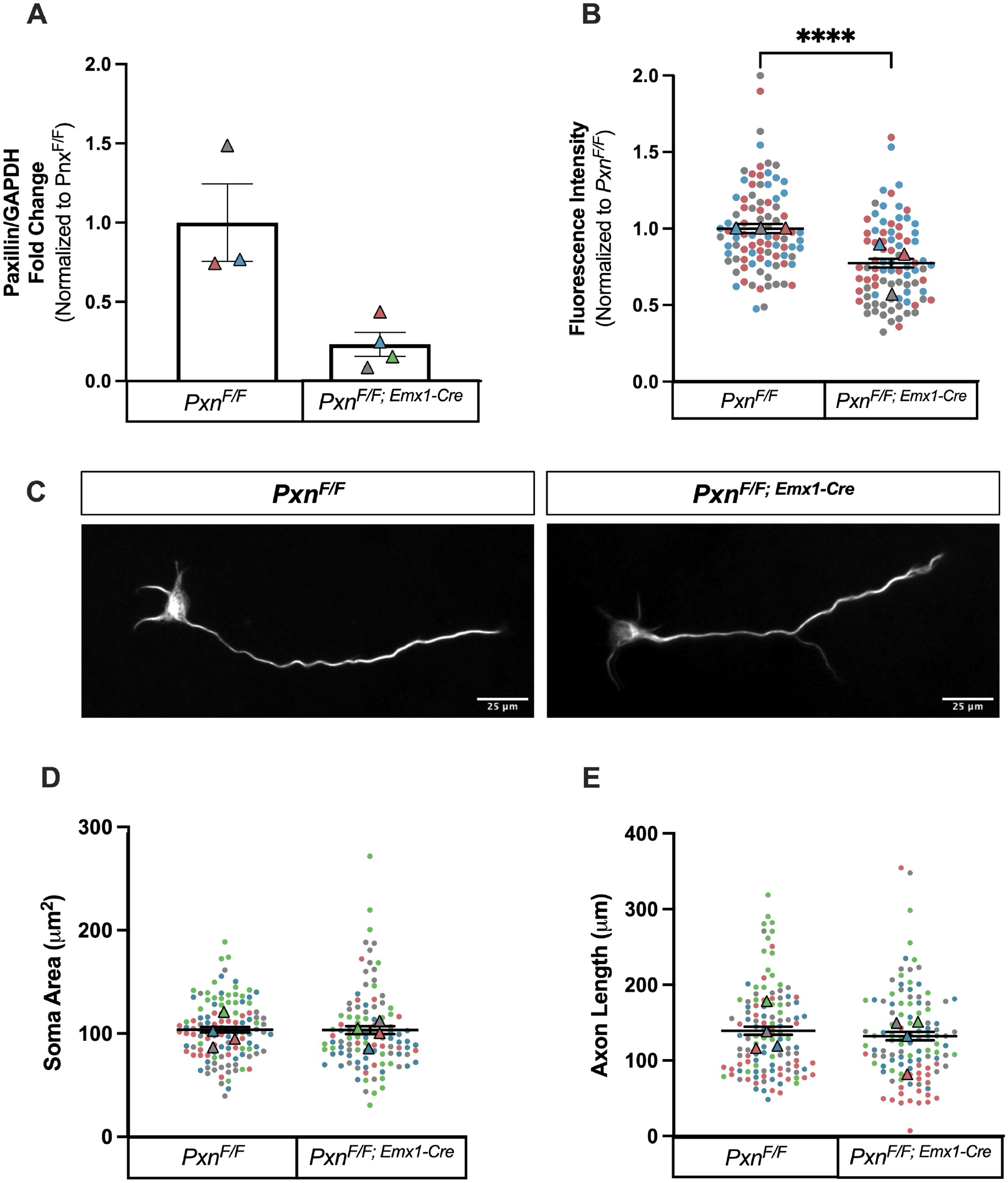
Cortical neurons cultured from *Pxn*^*F/F; Emx1-Cre*^ exhibit a significant decrease in paxillin expression, but this does not affect soma area or axon length. **A)** ddPCR of paxillin mRNA levels in *Pxn*^*F/F*^ and *Pxn*^*F/F; Emx1-Cre*^ neurons, normalized to GAPDH. Mann-Whitney, p = 0.0571. *Pxn*^*F/F*^, n=3 animals; *Pxn*^*F/F; Emx1-Cre*^, n=4 animals. **B)** Paxillin was quantified in exposure-matched images of paxillin immunostained neurons. Paxillin was quantified in the entire cell. Mann-Whitney, p < 0.0001. *Pxn*^*F/F*^, n=91 neurons; *Pxn*^*F/F; Emx1-Cre*^, n=86 neurons. This experiment was repeated using 3 individual animals. **C)** Representative exposure-matched images of *Pxn*^*F/F*^and *Pxn*^*F/F; Emx1-Cre*^ neurons immunostained for β-tubulin. Scale Bars, 25μm. **D-E)** Soma area and axon length were quantified in *Pxn*^*F/F*^ and *Pxn*^*F/F; Emx1-Cre*^ neurons. Mann-Whitney, p = 0.3164 for soma area, p = 0.4177 for axon length. Axon Length: *Pxn*^*F/F*^, n=114 neurons; *Pxn*^*F/F; Emx1-Cre*^, n=109 neurons. Soma area: *Pxn*^*F/F*^, n=110 neurons; *Pxn*^*F/F;* Emx1-Cre^, n=91 neurons. This experiment was repeated using 4 individual animals.

Paxillin is part of the adhesion complex that regulates migration^4^. Thus, we examined how paxillin loss affects developing neuron morphology. We cultured E17 cortical neurons from control (*Pxn*^*F/F*^) and conditional paxillin knockout *(Pxn*^*F/F; Emx1-Cre*^) animals for 2 days *in vitro* (DIV). Cells were fixed and imaged to quantify soma area and axon length under basal conditions (**Figure 1C**). Interestingly, there were no significant differences in soma area or axon length in neurons from *Pxn*^*F/F; Emx1-Cre*^ as compared to their control uterine mates (**Figure 1D-E**).

A recent study examining paxillin knockdown in chick motor neurons also reported no significant differences in axon length at basal conditions. However, paxillin knockdown did cause an axon guidance deficit in response to extracellular guidance cues involved in limb development^7^. Thus, we investigated whether morphological parameters may be altered in *Pxn*^*F/F; Emx1-Cre*^ neurons following stimulation with BDNF, which promotes axon growth. Cortical neurons cultured for 2 DIV were starved for 3 hours by removing B27 from the culture media. After the starvation period, B27 and 100 ng/ml BDNF were added to the culture media for 20 minutes, and then the neurons were fixed (**Figure 2A-C**)^8^. There was no significant change in soma area following BDNF stimulation (**Figure 2B**). However, there was an increase in axon length of *Pxn*^*F/F; Emx1-Cre*^ neurons under BDNF-stimulated conditions, as compared to control (**Figure 2C**).

**Figure 2.**
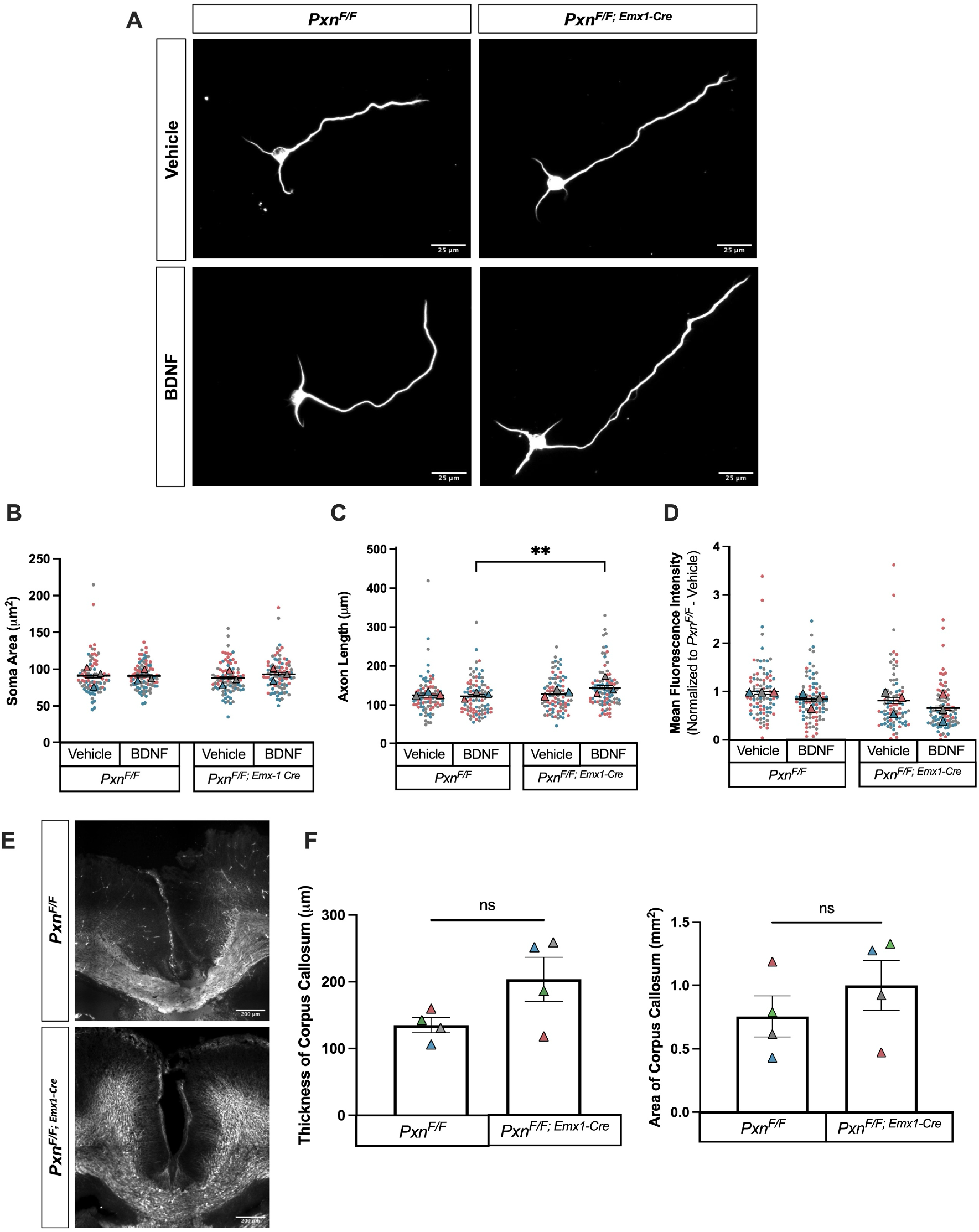
BDNF stimulated cortical neurons cultured from *Pxn*^*F/F; Emx1-Cre*^ mice have increased axon length, but no difference in soma area, β-tubulin expression, or thickness or area of the corpus callosum. **A)** Cortical neurons were cultured for 2DIV before being starved for 3 hours. Following starvation, neurons were stimulated with BDNF for 20 minutes. Representative exposure-matched images of *Pxn*^*F/F*^ and *Pxn*^*F/F; Emx1-Cre*^ cortical neurons immunostained for β-tubulin under vehicle or BDNF stimulated conditions. Scale bars, 25µm. **B)** Quantification of soma area in *Pxn*^*F/F*^ and *Pxn*^*F/F; Emx1-Cre*^ cortical neurons under vehicle or BDNF stimulated conditions. 2-way ANOVA. Interaction, p = 0.2437; Genotype, p = 0.9009; Treatment, p = 0.2686. *Pxn*^*F/F*^ + vehicle, n=87 neurons; *Pxn*^*F/F*^ + BDNF, n=90 neurons; *Pxn*^*F/F; Emx1-Cre*^ + vehicle, n=89 neurons, *Pxn*^*F/F; Emx1-Cre*^ + BDNF, n=90 neurons. **C)** Axon length of *Pxn*^*F/F*^ and *Pxn*^*F/F; Emx1-Cre*^ cortical neurons under vehicle or BDNF stimulated conditions. 2-way ANOVA. Interaction, p = 0.0637; Genotype, *p = 0.0133; Treatment, p = 0.1779. Šídák Post-Hoc: *Pxn*^*F/F*^ + vehicle vs. *Pxn*^*F/F*^ + BDNF, p = 0.9938; *Pxn*^*F/F*^ + vehicle vs *Pxn*^*F/F; Emx1-Cre*^ + vehicle, p = 0.9864; *Pxn*^*F/F*^ + BDNF vs *Pxn*^*F/F; Emx1-Cre*^ + BDNF, **p = 0.0085; *Pxn*^*F/F; Emx1- Cre*^ + vehicle vs *Pxn*^*F/F; Emx1-Cre*^ + BDNF, p = 0.0904. *Pxn*^*F/F*^ + vehicle, n=87 neurons; *Pxn*^*F/F*^ + BDNF, n=90 neurons; *Pxn*^*F/F; Emx1-Cre*^ + vehicle, n=89 neurons, *Pxn*^*F/F; Emx1-Cre*^ + BDNF, n=90 neurons. **D)** Fluorescence intensity of β-tubulin in the axon under vehicle or BDNF stimulated conditions. 2-way ANOVA. Interaction, p = 0.9566; Genotype, ***p = 0.001; Treatment, **p = 0.0038. Šídák Post-Hoc: *Pxn*^*F/F*^ + vehicle vs. *Pxn*^*F/F*^ + BDNF, p = 0.1376; *Pxn*^*F/F*^ + vehicle vs *Pxn*^*F/F; Emx1-Cre*^ + vehicle, p = 0.1866; *Pxn*^*F/F*^ + BDNF vs *Pxn*^*F/F; Emx1-Cre*^ + BDNF, p = 0.1806; *Pxn*^*F/F; Emx1-Cre*^ + vehicle vs *Pxn*^*F/F; Emx1-Cre*^ + BDNF, p = 0.1577. *Pxn*^*F/F*^+ vehicle, n=87 axons; *Pxn*^*F/F*^ + BDNF, n=89 axons; *Pxn*^*F/F; Emx1-Cre*^ + vehicle, n=87 axons, *Pxn*^*F/F; Emx1-Cre*^ + BDNF, n=88 axons. **E)** Representative images of P0 coronal brain sections from *Pxn*^*F/F*^ and *Pxn*^*F/F; Emx1-Cre*^ animals, showing the body of the corpus callosum. **F)** Quantification of thickness and area of the corpus callosum. Mann-Whitney, p = 0.2000 for corpus callosum thickness, p = 0.3429 for corpus callosum area. All experiments were repeated using 3-4 individual animals.

Because there was an increase in axon length in *Pxn*^*F/F; Emx1-Cre*^ mice under BDNF-stimulated conditions, we examined the expression of β-tubulin because it binds paxillin and is critical for axon outgrowth^4^. Thus, we quantified β-tubulin expression in axons, comparing paxillin knockout to control (**Figure 2C**). Although there was a main effect of genotype, β-tubulin expression in *Pxn*^*F/F; Emx1-Cre*^ axons under BDNF-stimulated conditions was not significantly altered as compared to control BDNF-stimulated neurons (**Figure 2D**).

To further investigate whether the loss of paxillin affects axon growth and guidance and validate the *in vitro* findings *in vivo*, the corpus callosum was examined in postnatal day 0 (P0) brains from *Pxn*^*F/F*^ and *Pxn*^*F/F; Emx1-Cre*^ animals. We focused on the corpus callosum because pyramidal neurons, which are targeted in the Emx1-Cre model and studied in the *in vitro* experiments, form this axon bundle. We measured the thickness and area of the axonal fibers in the middle of the corpus callosum ^9, 10^. There was a slight increase in corpus callosum thickness and area in *Pxn*^*F/F; Emx1-Cre*^ animals, but this was not significant (**Figure 2E-F**). Thus, loss of paxillin does not affect pyramidal neuron axons that form the corpus callosum.

Taken together, these findings suggest that paxillin may be dispensable for axon growth of cortical neurons. However, more research is needed, as previous studies suggest paxillin contributes to neurite initiation of other cell types grown on soft substrates^11^. It is also possible that although paxillin does not regulate axon growth, it may regulate axon guidance in response to specific guidance cues; this is suggested by a previous study^7^ and our BDNF stimulation data. More experiments are needed to rigorously assess and understand the BDNF-induced increase in axon length in paxillin knockout animals. Additionally, other members of the paxillin family of adapter proteins (i.e., Hic-5 or leupaxin) may compensate for the loss of paxillin in these neurons^12^. Further studies in *Pxn*^*F/F; Emx1-Cre*^ neurons will provide more insight into the mechanisms that may be regulating axon growth and guidance following the loss of paxillin.

## 3. MATERIALS AND METHODS

### 3.1 Animals and cell culture

All procedures were approved by the University of South Carolina IACUC. *Pxn*^*F/F*^ mice were obtained from the Laboratory of Dr. Turner at SUNY Upstate Medical University (Jackson Laboratory stock #035946, RRID:IMSR_JAX:035946)^2^. The Emx1-IRES-Cre mice were obtained from The Jackson Laboratory (Jackson Laboratory stock #005628, RRID:IMSR_JAX:005628)^6^. Animals were mated in a series of crosses to generate *Pxn*^*F/F*^ and *Pxn*^*F/F; Emx1-Cre*^ animals (see Graphical Abstract). Cortical neurons were extracted from timed pregnant mice on E17 and plated onto 100μg/mL poly-L-lysine (Sigma, Cat# P1274) and 10μg/mL laminin (Gibco, Cat# 23017015) coated coverslips as previously described^8^. A piece of cortex or tail clip from each animal was collected to extract DNA. All animals were genotyped using PCR prior to use, using primers as previously described^2, 6^.

### 3.2 Immunocytochemistry

Immunocytochemistry experiments were performed as previously described^8^. The following primary antibodies were used: rabbit anti-paxillin (1:500; Abcam Cat# Ab32084, RRID:AB_779033) and mouse anti-β-tubulin (1:1,000; DSHB Cat# E7, RRID:AB_528499). The following secondary antibodies were used: donkey anti-rabbit Alexa 488 (1:1,000; Life Technologies) and goat anti-mouse Alexa 568 (1:1,000; Life Technologies).

### 3.3 Digital Droplet PCR

RNA from cortical neurons was lysed and extracted using a RNeasy Plus Kit (Qiagen). Reverse transcription used the SensiFast™ cDNA synthesis kit (Bioline, BIO-65053). ddPCR products were quantified using the QX200™ ddPCR system and EvaGreen Supermix (Biorad). Paxillin levels of *Pxn*^*F/F*^ and *Pxn*^*F/F; Emx1-Cre*^ neurons were normalized to GAPDH. Primer sets for paxillin (Mm.PT.58.8475697) and GAPDH (Mm.PT.39a.1) were purchased from IDT.

### 3.4 Immunohistochemistry (IHC)

For IHC experiments, postnatal day 0 brains were fixed in 4% paraformaldehyde for 6 hours at 4°C as previously described ^9^. Coronal sections of 30 μm were obtained using a cryostat. Slides were washed with 1X PBS, then blocked in 5% goat serum. The primary antibody rat anti-L1 (1:500; Millipore Cat# MAB5272, RRID:AB_2133200) was incubated overnight at 4°C. The secondary antibody Donkey Anti-Rat IgG (1:500; Jackson ImmunoResearch Labs Cat# 712-165-153, RRID:AB_2340667) was incubated for 1 hour at room temperature.

### 3.5 Image Acquisition and Analysis

Images were obtained on a Nikon Ti2-E microscope, keeping all acquisition parameters consistent across experiments. Quantifications were made using Fiji. Axon length is the length of the longest neurite from the cell body to the tip of the growth cone. The thickness and area of the corpus callosum were measured as previously described^9, 10^. The thickness is measured down the middle of the fiber bundle. The area outlined the middle fiber bundle before it curves up into each hemisphere.

### 3.6 Statistical Analysis

Statistical analyses were performed in GraphPad Prism with a p-value set at <0.05. Data is presented as mean + SEM.

## AUTHOR CONTRIBUTIONS

Katelyn Rygel designed the study, conducted experiments and analyzed the data, maintained animals, and wrote the original draft of the manuscript. Kasey Gillespie conducted experiments and analyzed the data. Kristy Welshhans designed the study, reviewed and edited the manuscript, obtained funding, and supervised the study. All authors reviewed and edited the manuscript before submission.

## ACKNOWLEDGMENTS

We thank Christopher Turner for generously donating the paxillin floxed mice to us. The graphical abstract was made in BioRender.com. Research reported in this publication was supported by the University of South Carolina School of Medicine Instrumentation Resource Core Facility (RRID:SCR_024955). The content is solely the responsibility of the authors and does not necessarily represent the official views of the National Institutes of Health, Columbia VA Healthcare System, or the USC Vice President of Research Office.

## DATA AVAILABILITY STATEMENT

The raw data supporting this study’s findings are available from the corresponding author upon request.

## REFERENCES

1. Nichol, R. I., Hagen, K. M., Lumbard, D. C., Dent, E. W., and Gomez, T. M. (2016) Guidance of Axons by Local Coupling of Retrograde Flow to Point Contact Adhesions. J Neurosci 36, 2267–2282

2. Rashid, M., Belmont, J., Carpenter, D., Turner, C. E., and Olson, E. C. (2017) Neural-specific deletion of the focal adhesion adaptor protein paxillin slows migration speed and delays cortical layer formation. Development 144, 4002–4014

3. Baas, P. W., Rao, A. N., Matamoros, A. J., and Leo, L. (2016) Stability properties of neuronal microtubules. Cytoskeleton (Hoboken) 73, 442–460

4. Efimov, A., Schiefermeier, N., Grigoriev, I., Ohi, R., Brown, M. C., Turner, C. E., Small, J. V., and Kaverina, I. (2008) Paxillin-dependent stimulation of microtubule catastrophes at focal adhesion sites. J Cell Sci 121, 196–204

5. Hagel, M., George, E. L., Kim, A., Tamimi, R., Opitz, S. L., Turner, C. E., Imamoto, A., and Thomas, S. M. (2002) The adaptor protein paxillin is essential for normal development in the mouse and is a critical transducer of fibronectin signaling. Mol Cell Biol 22, 901–915

6. Gorski, J. A., Talley, T., Qiu, M., Puelles, L., Rubenstein, J. L. R., and Jones, K. R. (2002) Cortical Excitatory Neurons and Glia, But Not GABAergic Neurons, Are Produced in the Emx1-Expressing Lineage. The Journal of Neuroscience, 6309–6314

7. Tsai, W. L., Chang, C. J., Wang, C. Y., Hsu, T. I., Chang, M. Y., Wu, Y. H., Chang, P. S., Lin, K. L., Chuang, J. Y., Kania, A., and Kao, T. J. (2021) Paxillin Is Required for Proper Spinal Motor Axon Growth into the Limb. J Neurosci 41, 3808–3821

8. Kershner, L., and Welshhans, K. (2017) RACK1 is necessary for the formation of point contacts and regulates axon growth. Dev Neurobiol 77, 1038–1056

9. Jain, S., Watts, C. A., Chung, W. C. J., and Welshhans, K. (2020) Neurodevelopmental wiring deficits in the Ts65Dn mouse model of Down syndrome. Neurosci Lett 714, 134569

10. Schreiber, J., Grimbergen, L. A., Overwater, I., Vaart, T. V., Stedehouder, J., Schuhmacher, A. J., Guerra, C., Kushner, S. A., Jaarsma, D., and Elgersma, Y. (2017) Mechanisms underlying cognitive deficits in a mouse model for Costello Syndrome are distinct from other RASopathy mouse models. Sci Rep 7, 1256

11. Chang, T. Y., Chen, C., Lee, M., Chang, Y. C., Lu, C. H., Lu, S. T., Wang, D. Y., Wang, A., Guo, C. L., and Cheng, P. L. (2017) Paxillin facilitates timely neurite initiation on soft-substrate environments by interacting with the endocytic machinery. Elife 6

12. Brown, M. C., and Turner, C. E. (2004) Paxillin: adapting to change. Physiological Reviews 84, 1315–1339

